# Human subsidies facilitate hyperpredation of Mediterranean island wildlife by outdoor cats

**DOI:** 10.1101/2025.07.27.667076

**Authors:** Jeffrey R. Ferrer, John Vandermeer, Collin J. Richter, Erin R. Baldwin, Johannes Foufopoulos

**Author notes:** Corresponding Author (JRF). Collin J. Richter, John Vandermeer, and Erin R. Baldwin contributed equally to this work.

## Abstract

Domestic cats (*Felis catus*) are avid wildlife predators and one of the most harmful invasive species. The lethal impacts of an introduced predator such as cats on wildlife can be further exacerbated by the introduction of an additional abundant non-native prey species capable of supporting an exceptionally dense predator population, a phenomenon known as hyperpredation. A special case of hyperpredation involves human food subsidies, when an invasive predator impacting native wildlife is supported not by another invasive taxon, but by human-derived food sources. To test whether access to anthropogenic food subsidies by cats is causing hyperpredation, mark-recapture methodology was used to measure twelve cat populations experiencing a gradient of human subsidies on the Mediterranean island of Naxos, Greece. Line-transect surveys were conducted at each site to measure reptile population abundance across this gradient, and other factors including human density and distance of reptiles from villages were considered as well. Strong evidence was found that the population size of cats is a direct result of the density of anthropogenic food available to them, and a corresponding decline in the size of reptile populations as cat density increased was observed. It was also shown that as cat populations exceed the available supply of human food, the negative effects of cat density on reptile populations are exacerbated. These results demonstrate that access to anthropogenic subsidies has allowed cat populations to expand to exceptional levels, driving hyperpredation on island wildlife.

## Introduction

Invasive species are one of the greatest threats to biodiversity worldwide [1–3]. Invasive species have been implicated in over 50% of all modern extinctions whose causes are known, and in about 20% of total cases, invasive species have been identified as the sole driver of extinction [4–5]. One of the most harmful groups of invasive species are mammals, who currently threaten more IUCN listed critically endangered terrestrial vertebrate species than any other group, an impact driven primarily by black rats (*Rattus rattus*) and domestic cats (*Felis catus*) [6].

Cats have been linked to ≥ 63 vertebrate extinctions [7] and are recognized as one of the most harmful invasive species due to their global distribution and devastating impacts on wildlife [8]. In the United States alone, it has been estimated that cats kill billions of birds, mammals, and reptiles each year, with mortality driven primarily by unowned free-roaming cats, rather than pets [9]. In addition to direct predation, cats can have many other negative effects on wildlife such as influencing prey behavior by increasing prey wariness and decreasing foraging time [10–11], transmission of diseases such as rabies, toxoplasmosis, cutaneous larval migrans, tularemia, and plague [12], and loss of genetic integrity in native species such as the Eurasian wildcat (*Felis silvestris silvestris*) through hybridization [13].

The damaging effects of cats are particularly prevalent on islands [14–15]. Despite making up only 5.3% of the Earth’s landmass, 61% of all known extinctions have occurred on islands, and invasive species have been cited as the most frequent cause of insular species extinctions [16]. Island communities are disproportionately affected by invasive mammalian predators, and especially cats, where a general lack of native predators typically results in down-regulated antipredator responses, rendering island wildlife relatively tame and easy to capture [17–19]. Cats have emerged as the causal factor in at least 33 extinctions of insular vertebrate species [9], and because they have already been introduced to most suitable islands worldwide [14], they represent a threat to numerous additional island endemics [10].

The impacts of an introduced predator on native prey can be exacerbated by the introduction of an additional non-native prey species, a phenomenon known as hyperpredation [20–23]. When this occurs, a rapidly reproducing non-native prey species supports an unusually dense population of predators. This can have severe consequences for the native prey species, which may be decimated by the resulting increase in predators. For example, the extinction of the endemic Macquarie Island Parakeet (*Cyanoramphus novaezelandiae erythrotis*) is thought to have occurred following hyperpredation by cats. Although cats were introduced onto Macquarie Island within ten years of its discovery in 1810, the native parakeet withstood predation pressure and remained numerous there until about 1880. In 1879, European rabbits (*Oryctolagus cuniculus*) were released onto the island and reproduced rapidly. Within a few years, cat populations persisting primarily on rabbits expanded rapidly and opportunistically consumed parakeets. The parakeet was last definitively seen in 1890 [18, 20, 24]. Hyperpredation has since been recorded in several systems [25–28], and understanding how to predict and manage it will only increase in importance as the spread of invasive species is expected to increase with the expansion of global trade networks [29].

A related situation can occur when an invasive predator, impacting native wildlife, is supported not by another invasive taxon, but by human-derived food sources [30] (Fig 1). In many urban environments, densities of predators increase simply through ease of access to anthropogenic resources like trash and hand-outs [31–33]. For example, while cats are typically solitary and territorial hunters, the presence of large quantities of a stable food resource can alter their behavior by reducing their home-range size and territoriality, increasing tolerance of home-range overlap, and leading to the formation of cat colonies [34–35]. While densities of cats can vary greatly, Liberg et al. (2000) [31] found that densities above 100 cats/km^2^ are only found in urban areas with a constant source of anthropogenic food supplementation. These unnaturally high densities of cats can potentially result in a unique form of hyperpredation where the introduced ‘prey’ that is supplementing the predator population is simply anthropogenically derived food. As cats have been shown to have high fidelity to a single feeding site [34], it is expected that the number of feeding sites available to cats should be correlated with the size of the cat population. However, this has yet to be demonstrated, and the extent to which increased food supplementation affects greater impacts on wildlife remains unclear as for example, more subsidies may simply result in a switch away from wildlife to anthropogenic sources.

**Fig 1.**
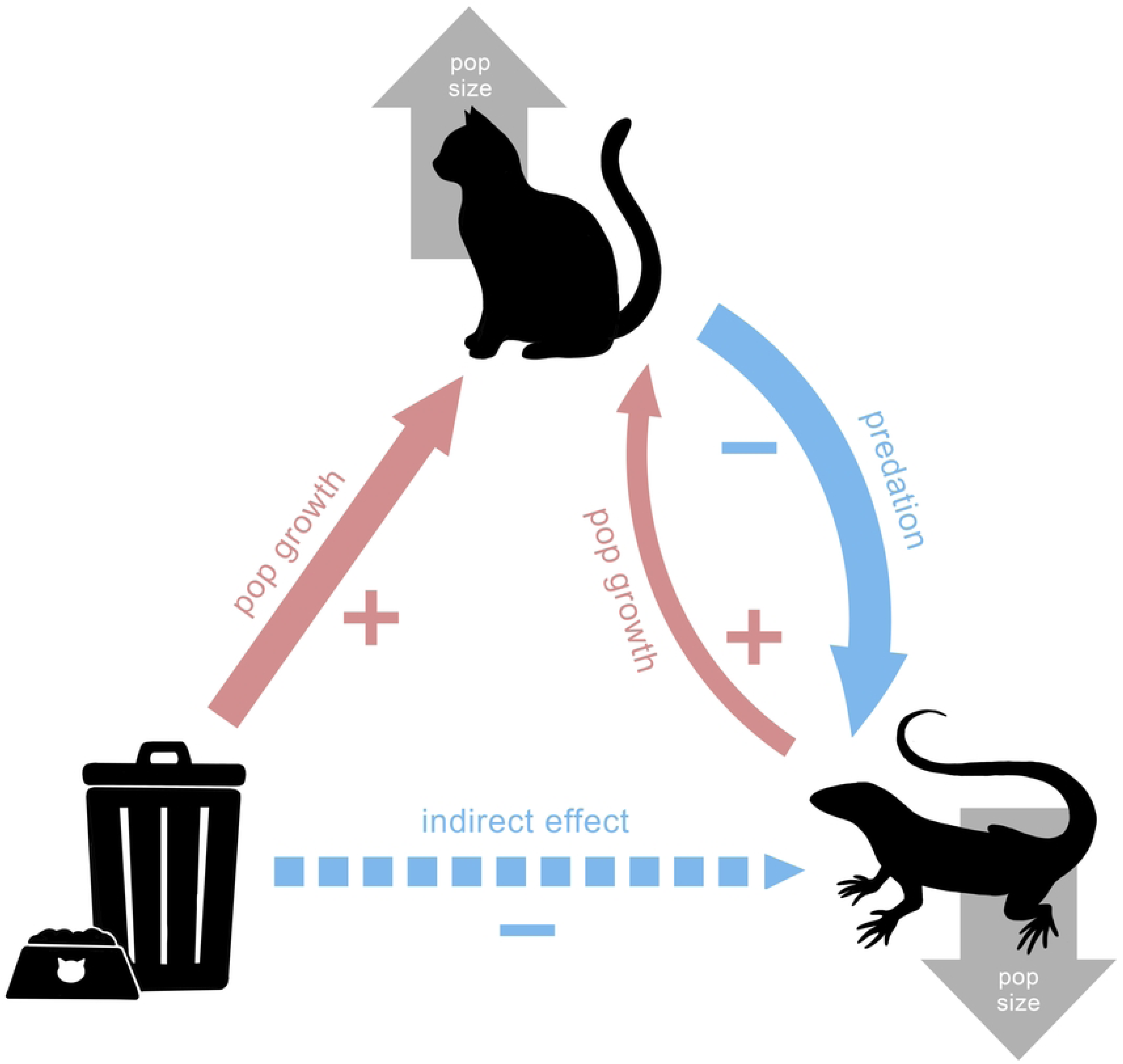
Conceptual diagram showing how the addition of anthropogenic food supplementation into a predatory-prey relationship can result in hyperpredation on wildlife.

Cats were first introduced to the Mediterranean islands thousands of years ago [36–38] and are playing an important predator role in both urban and rural Mediterranean ecosystems [39–40] where they have been implicated in significant wildlife mortality [11,41,42]. The goals of this study were to establish which factors support cat populations in a representative Mediterranean ecosystem (the island of Naxos in the Aegean Sea, Greece), and to determine how cats are affecting resident wildlife populations while considering other factors that may shape a possible relationship between cats and wildlife.

Methods used to answer these questions included, 1) obtaining population and density estimates of twelve discrete cat populations experiencing a range of food supplementation, 2) quantifying the amount of food supplementation available to each cat population, 3) surveying reptile communities in the natural habitat in the immediate vicinity of each of these locations, and 4) providing visual support through the development of heat maps showing the spatial distribution of cats.

By investigating these population level relationships along a gradient of food supplementation within a representative island system, this study is taking a novel and fine-grained approach that will allow a quantitative demonstration of the connection between human food subsidies and decreases in wildlife in Mediterranean wildlife.

## Methods

### Study Site

Field work was conducted during the months of May and June 2024 on the Aegean Island of Naxos, Greece. Naxos (430km^2^) is the largest island in the Cyclades archipelago with a typical Mediterranean climate consisting of warm dry summers, and mild moist winters. The landscape of Naxos is characterized by xeric *phrygana*, a type of thorny summer-deciduous Mediterranean cushion scrub community dominated by *Sarcopoterium spinosum* and *Genista acanthoclada*, as well as high evergreen *maquis* dominated by *Quercus coccifera* and *Juniperus turbinata*. The wide distribution of these communities is consistent with a long history of disturbance and degradation of the original oak forest habitats through grazing and agricultural activities on the island [43]. Areas near villages consist of a mosaic of these communities mixed with irrigated gardens, grainfields, as well as olive groves.

Cycladic wildlife is dominated by reptiles which, by virtue of their life history, are particularly well suited for the Mediterranean climate [44]. Naxos is home to 13 reptile species, including two introduced species (*Hemidactylus turcicus* and *Chalcides ocellatus*). While reptiles are overall common, communities on the island are dominated by the Aegean wall lizard (*Podarcis erhardii*), hereafter “wall lizard”, which can easily be observed. Both wall lizards and other reptile taxa occur in particularly high densities along the drystone walls and terraces which act as focal points of reptile activity, and are a ubiquitous feature of the Mediterranean landscape [45–46]. Reptile populations in the region are likely shaped by a combination of both bottom-up, as well as top-down effects. Among lizard predators, snakes are probably the most important [47]. The only other mammalian predator besides cats are stone martens (*Martes foina*), although because of their strictly nocturnal activity pattern, they do not likely encounter the broadly heliothermic reptiles. Because of this high aggregate biomass, reptiles are a particularly important component of Cycladic ecosystems and constitute an ideal system to investigate the impacts of cats on local species communities.

Naxos contains over 30 towns and villages that range in population size from being essentially abandoned, to the capital of Chora (with ca. 6000 inhabitants) [48]. Naxian settlements have traditionally been built in a very compact manner, facilitating defense against pirate raids [49]. They typically encompass a dense network of stone-lined footpaths that allow access to every part of the village. Household refuse is collected in select sites located at the edge of a settlement where garbage truck access is possible, creating distinct hot spots of waste accumulation. At the same time, there is a clear border and very sharp transition from the last outer row of houses to the surrounding agricultural matrix where resident wildlife occurs.

Cats can be found in large numbers in nearly every town on Naxos. At the same time, because of Naxos’ arid landscape, cats are present but rare in the remote countryside, especially more than 1-2km away from human habitations [11]. They appear to be highly dependent on the shelter of human settlements where they readily consume nutritionally dense cat food provided by residents as well as garbage from open dumpsters (S1 Appendix). These conditions are representative of many coastal areas around the Mediterranean Basin [40]. Within villages, cats exist across a spectrum of associations with humans, ranging from fully domesticated and fed to completely ignored and even persecuted (personal observation). Even when affiliated with a household, cats on Aegean islands are traditionally kept as outdoor pets and are allowed to roam freely. Because they are typically fed in a yard or in the street, and because of the open nature and dense path network of island villages, it is possible to obtain a good estimate of the extent of food supplementation in each village.

This study focuses on 12 representative villages selected randomly as study sites (Fig 2, S1 Table) while ensuring that the following criteria were met: 1) presence of a discrete village edge and a sufficiently dense network of village paths allowing easy access and high visibility to the entire settlement, and 2) the presence of dry-stone walls extending radially away from the village into the surrounding habitats to facilitate reptile surveys. Two of these rock walls ( ≥ 200m in length) were randomly selected for surveys for each village, except for one smaller village that contained a single wall.

**Fig 2.**
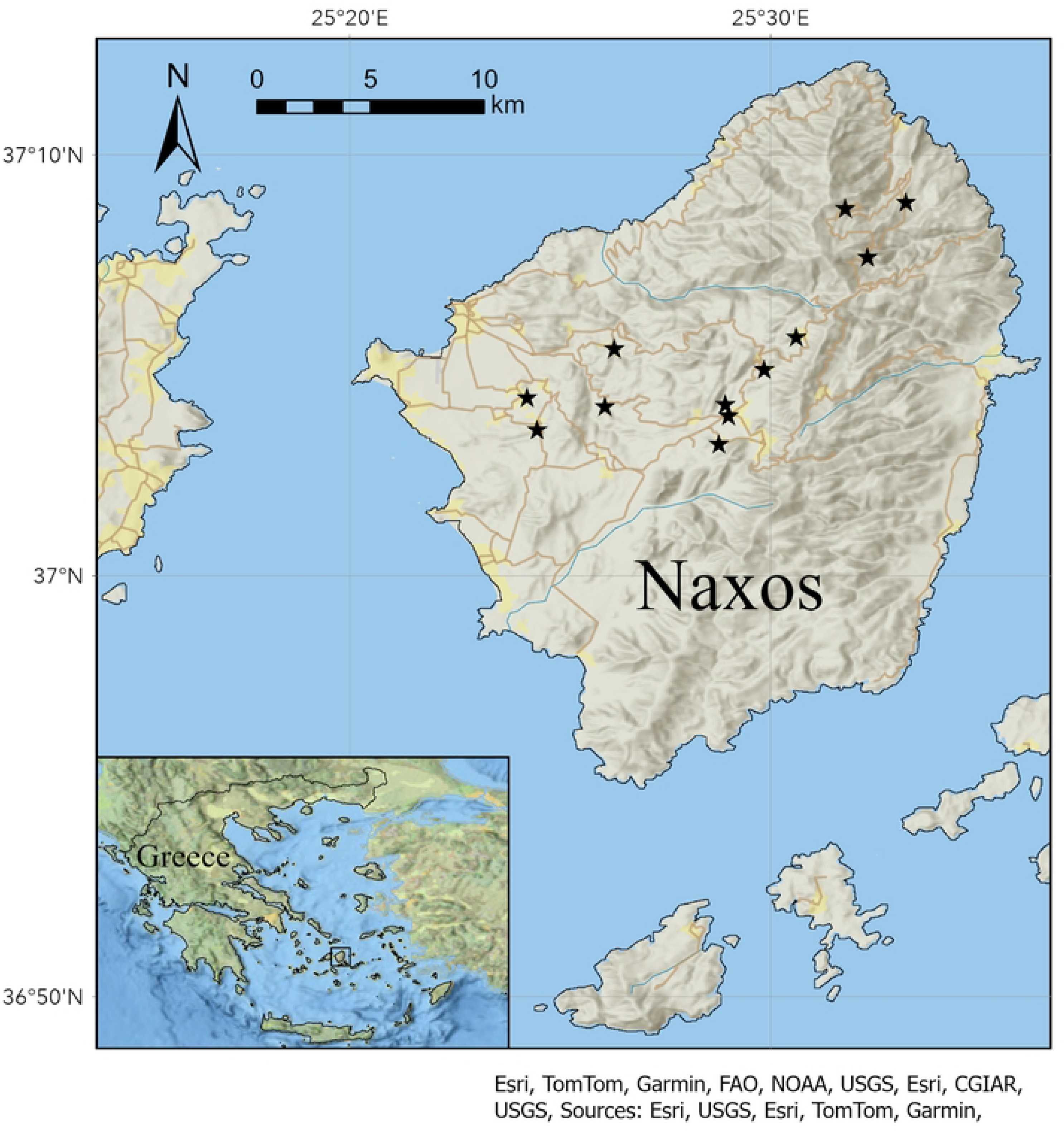
Map of the study area. Survey locations are indicated by black stars. Naxos, Greece and surrounding islets shown in main map. Entire country outlined in the inset, with a square around Naxos for reference) Map created in ArcGIS Pro v3.4 (2024). Produced with custom ESRI National Geographic Basemap (2025) [50–51].

### Reptile surveys

Reptile population abundance for each village were quantified through standardized surveys. Each survey was conducted along 200-meter transects parallel to drystone walls, which have been shown to act as foci for reptile activity in the region [11,52]. All surveys were completed during the months of May and June 2024 during peak reptile activity time (8am – 12pm), and under favorable weather conditions [53]. Weather metrics (temperature, wind speed, relative humidity, cloud cover) were collected at the beginning and end of each survey (Kestrel 5000 Environmental Meter, 2024, Nielsen-Kellerman Company, Boothwyn, Pennsylvania). Surveys began at the edge of a village and were conducted by slowly walking along the length of the drystone wall. Species name and distance (m) from the origin were recorded (Garmin, 2024, GPSMAP® 65s Handheld GPS device, Olathe, Kansas) for all reptiles observed on the wall or on the ground within 1 meter from the wall (using ‘unknown’ if no species ID was possible). Each wall received four surveys spread over the duration of the field season.

### Cat surveys

Cat surveys were conducted for three consecutive nights within each village. Surveys were performed in the evening during the time of peak cat activity (5pm – 8pm) and under appropriate weather conditions [54]. Surveys were conducted by walking along every path in a village once, and taking a picture and recording the GPS location of every cat observed. Survey tracks were recorded and uploaded into ArcGis Pro v3.4 [50] to confirm comprehensive coverage of each village. Great individual variability in terms of cat color, size, condition, and unique markings allowed cats to be identified to the level of individual [55]. Each cat was given a unique identifier code, and on the second and third survey nights, cats were designated as new individuals or recaptures from previous nights. Cat population size in each village was calculated using the Schnabel Index, a method used to estimate population size in mark-recapture studies when multiple sampling events are conducted [56–57]. Kittens were also recorded in the surveys but were excluded from the population size calculation due to their secretive nature making comprehensive identification difficult.

### Additional field data

During the first cat survey conducted at each village, the locations of all food available to cats in the form of either pet food dishes or communal garbage dumpsters were recorded. The aggregate number of food dishes and dumpsters were combined into a number of ‘food stations.’ Human population data for every community on the island were provided by the municipality of Naxos and confirmed through a review of the Greek National Statistical Agency. The area of each village (ha) was quantified in ArcGIS Pro v.3.4 by manually digitizing the perimeter of each village. Larger villages have more cats, as well as humans and feeding sites, necessitating the need to establish a correction for community size. To correct for area effects, cat population, number of food stations, and human population were divided by the village surface area.

### Statistical Analyses

Statistical analyses were performed in R version 4.4.2 [58] with the following the lme4 [59] and AICcmodavg [60] packages with the significance threshold set at 0.05.

### Cat density

Ordinary least squares regression (OLS) was used to first explore univariate associations between cat density, human density, and food station density and then to develop more complex models that included combinations of these predictors and their interactions. Corrected Akaike information criterion (AICc) [61] was used to determine the best fit model.

### Reptile populations

A cat ‘satiation index’ was calculated for each village as the residuals of the regression of cat density against food density. Positive residuals were interpreted as cats being in a food deficit, and therefore hungrier, while negative residuals were interpreted as cats being in a food surplus and more satiated. The residual values were rescaled so that the most satiated (index = 0) had a value of 0, and higher positive values indicated lower satiation, or increased hunger. This index was used as a model parameter to investigate whether level of cat satiation influenced reptile populations.

To analyze the overall effect of distance from a village on reptile populations, all reptile observations were combined, and the total number of reptiles observed at each 1-meter interval on the transects were calculated. These data met the assumptions of a linear regression, with a linear relationship between count and distance, normally distributed residuals, and homoscedasticity. Therefore, OLS was performed to analyze the association between distance and reptile counts for the combined dataset. Surveys for the village that only had one rock wall were given twice as much weight to prevent reptile counts for this village to be smaller due to reduced sampling effort. To further test whether cat density influenced the effect of distance on reptile counts, the number of reptiles observed at each 1-meter interval were summed within each village, and a Poisson regression was run for each village with reptile counts regressed against distance. The coefficient for distance was exponentiated to obtain a slope, and the slopes were regressed against cat density to investigate the association between cat density and the change in reptile counts with distance.

The association between reptile counts and cat density, cat satiation, and distance were analyzed in two separate analyses. For analysis one, reptile counts in each village were grouped into eight 25-meter distance bins. Then, through Poisson generalized linear mixed effect models (GLMM), with village ID included as a random effect, combinations of model parameters were explored to determine the best fit model, with AICc criteria used to determine model fit. An interaction term between cat density and satiation index was included in model exploration as well. For analysis two, reptile counts were analyzed at the survey level, through Poisson generalized linear mixed effect models (GLMM) with village ID included as a random effect. These models were used to analyze the association between the number of reptiles observed in each survey and cat density and cat satiation index across all villages without distance. An interaction term between cat density and satiation index was explored in this analysis as well.

## Results

### Cat density analysis

Cat densities in all villages were higher than what is normal for the species under rural or natural conditions (Fig 3, S2 Appendix) [31], except for one village with no cats that also had no additional food supplementation or permanent human residents.

**Fig 3.**
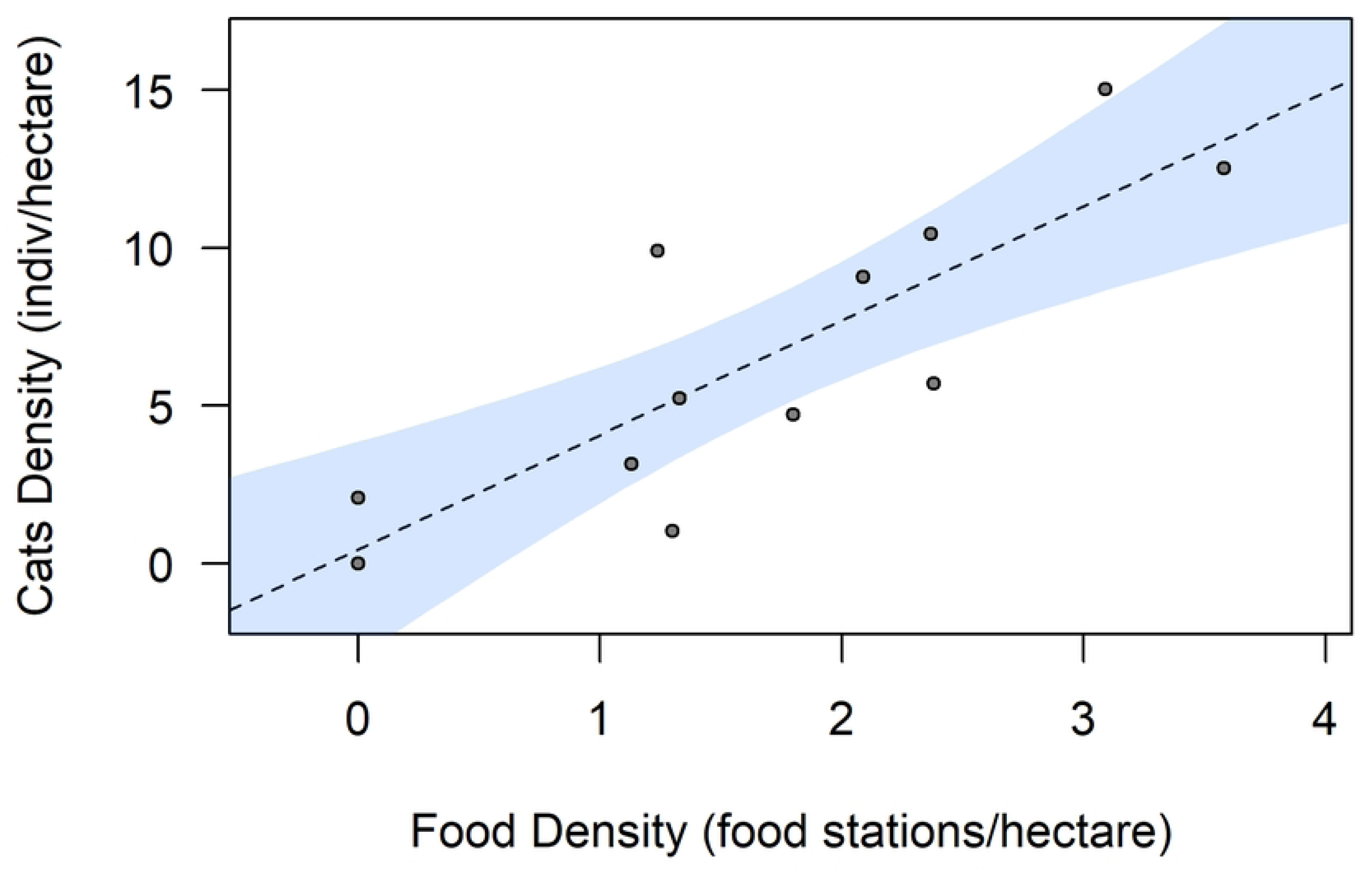
Ordinary least squares regression showing association between cat density and feeding density (p = 0.0008, adjusted R^2^ = 0.66). Shaded region represents 95% confidence interval.

Cat density was significantly positively associated with both predictor variables when analyzed independently-food density (adjusted R^2^ = 0.66, p = 0.009, n = 12, OLS, Fig 3), and human density (adjusted R^2^ = 0.45, p = 0.001, n = 12, OLS, Fig 4). When multiple predictor variables were included in model selection, two models showed nearly equal fit for predicting cat density. The first model (Model A) included only food density as a predictor (AICc = 65.54, AICcWt = 0.67, Log Likelihood = -28.27, Table 1) and the second model (Model B) included both food density and human density (AICc = 67.47, AICcWt = 0.26, Log Likelihood = -26.88, Table 1). Food density was significantly positively associated with cat density in both models (adjusted R^2^ = 0.657, p = 0.0001 for Model A and adjusted R^2^ = 0.698, p = 0.014 for Model B, Multiple Regression, Table 2), while human density was not significant in the model that included it. When the type of feeding supplementation (garbage bins and food dishes) was separated, cat density was significantly positively associated with both density of garbage bins (adjusted R^2^ = 0.28, p = 0.044, OLS) and density of feeding dishes (adjusted R^2^ = 0.73, p = 0.0002, OLS).

**Fig 4.**
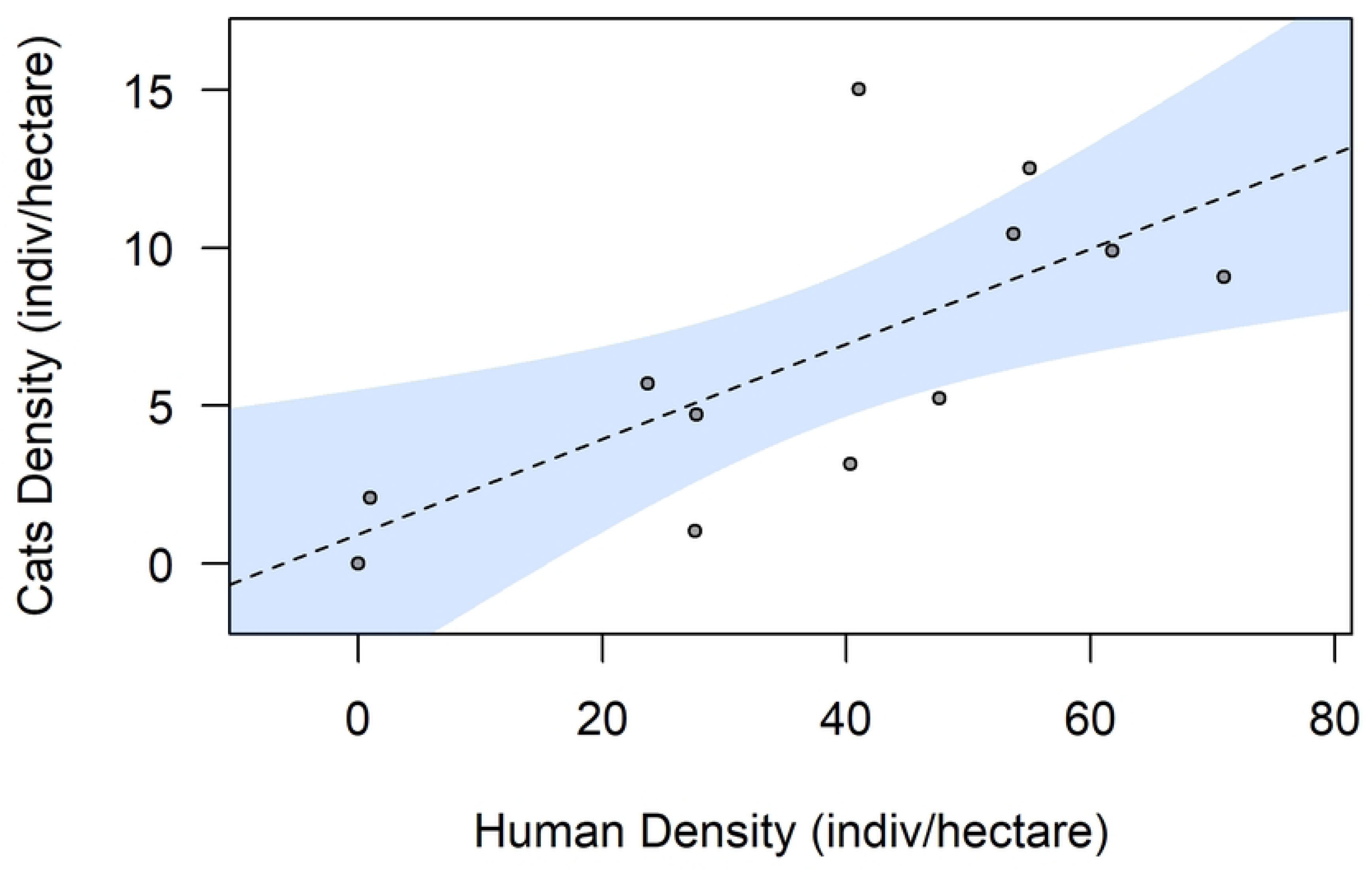
Ordinary least squares regression showing association between cat density and human density (p = 0.01, adjusted R^2^ = 0.45). Shaded region represents 95% confidence interval.

**Table 1:** Model selection for factors predicting density of cats, Models are ranked in ascending order by corrected Akaike corrected information criteria (AICc). ΔAICc, AICc weight, and Log Likelihood are provided for each model. Bold selections show models with near equal fit. Model variables include food density, human density, and an interaction between food.

| Predictors | AICc | ΔAICc | AICcWt. | Log Likelihood |
| --- | --- | --- | --- | --- |
| *Cat Density* |  |  |  |  |
| ~**food density** | **65.54** | **0.00** | **0.67** | **-28.27** |
| ~**human density + food density** | **67.47** | **1.93** | **0.26** | **-26.88** |
| ~human density | 71.2 | 5.66 | 0.04 | -31.10 |
| ~ human density + food density + interaction | 72.16 | 6.62 | 0.02 | -26.08 |
| null model | 75.87 | 10.33 | 0.00 | -35.27 |

**Table 2:**
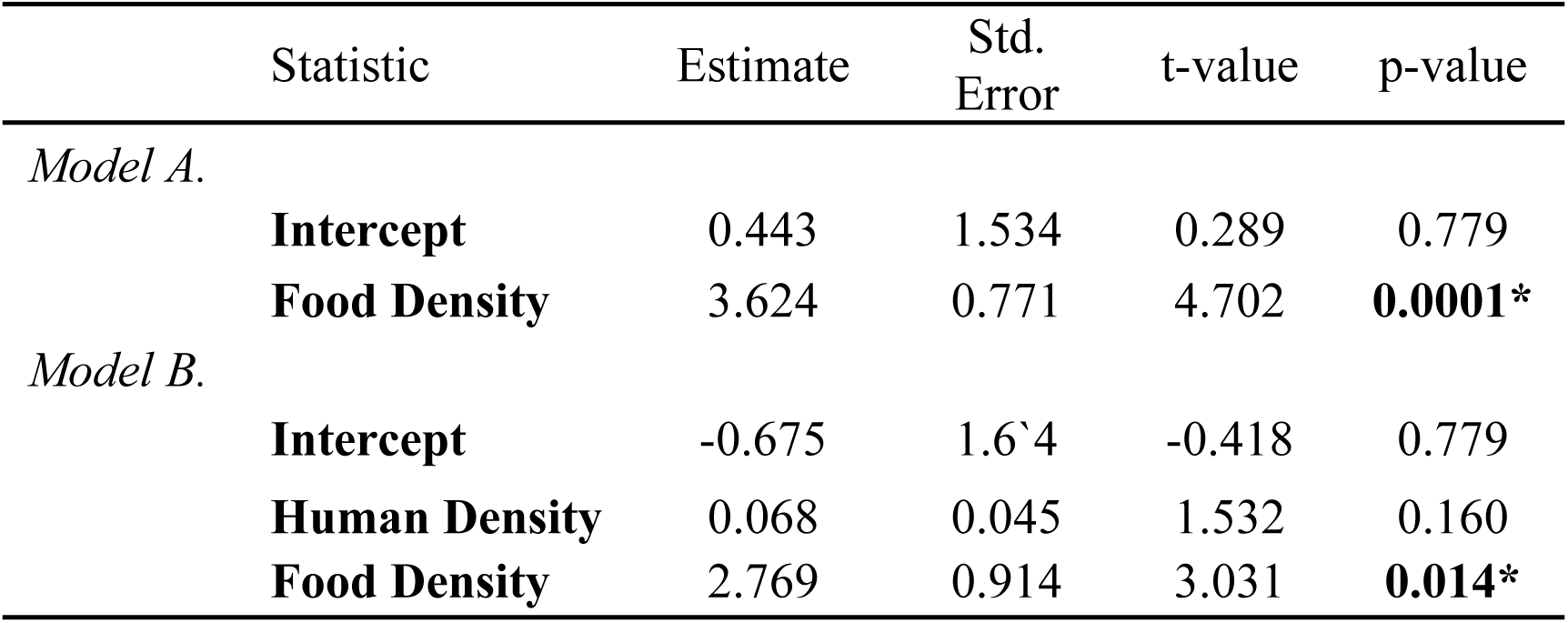
Results for cat density analysis using multiple linear regression showing two models with similar fit for predicting cat density. Asterisks indicate significant predictors (p < 0.05). Estimates, standard errors, and z-values are also shown. Model A adjusted R^2^ = 0.657. Model B adjusted R^2^ = 0.698.

### Reptile population analysis

A significant positive correlation was found between reptile counts and distance from the village edge, indicating that reptile numbers declined closer to human habitations and cats (Pearson’s R = .2176, p = 0.002, OLS, n = 200, Fig 5).

**Fig 5.**
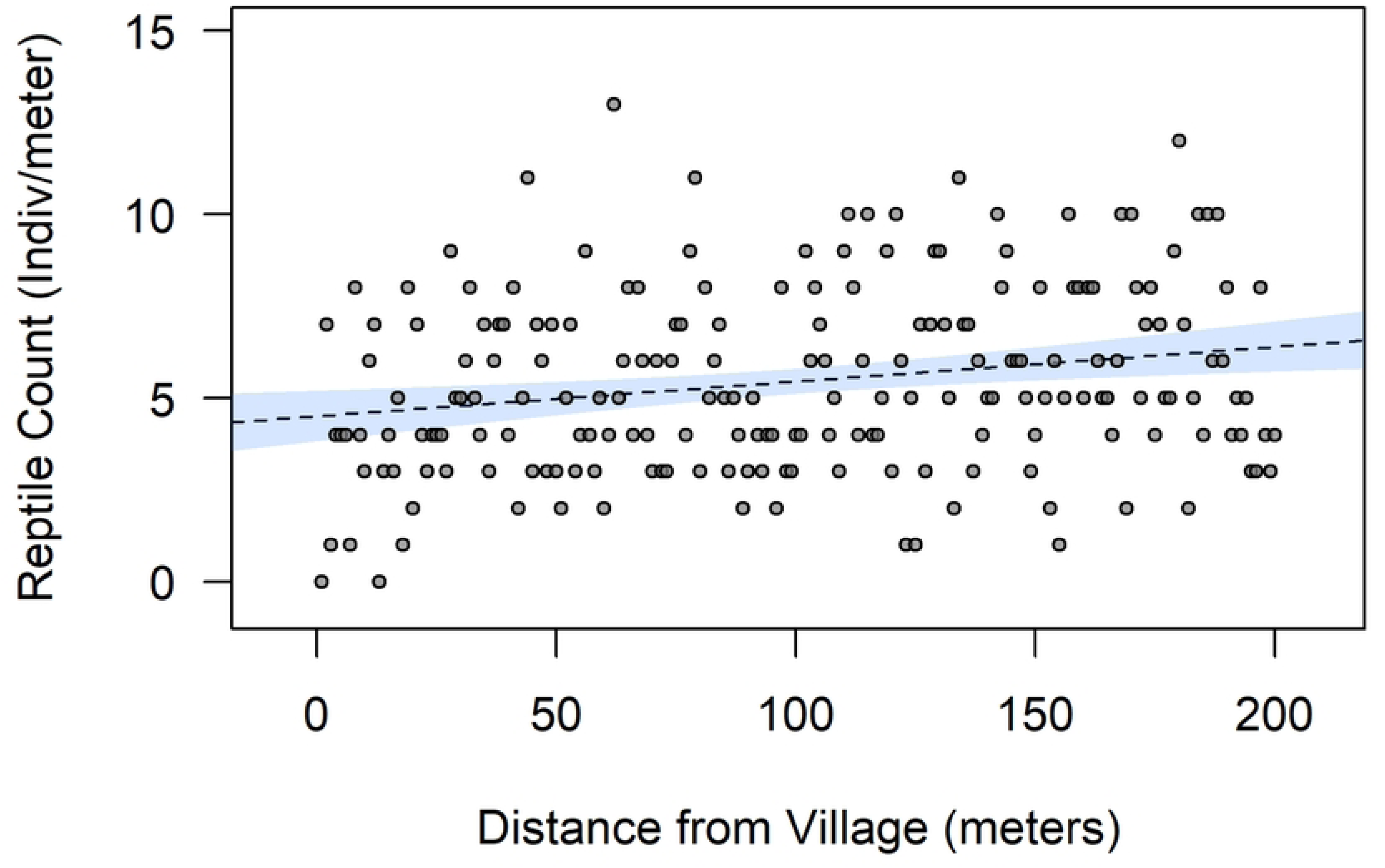
Ordinary least squares regression showing association between the number of reptiles observed at each 1-meter interval distance from village for the entire study (p = 0.002, adjusted R^2^ = 0.43). Shaded region represents 95% confidence interval.

For analysis one, when variables with the binned dataset were examined individually, Poisson GLMM’s showed that both cat density (marginal R^2^ = 0.33, conditional R^2^ = 0.82, p = 0.007, n = 96, Poisson GLMM), and satiation index (marginal R^2^ = 0.38, conditional R^2^ = 0.82, p = 0.002, n = 96, Poisson GLMM) were significantly negatively associated with reptile counts. When models that included different combinations of predictor variables and their interactions were compared, the model that best fit the data included all of the predictor variables; distance (p = 0.0003), cat density (p = 0.053), satiation index (p = 0.212), as well as the interaction between cat density and satiation index (p = 0.001) (marginal R^2^ = 0.67, conditional R^2^ = 0.83, AICc = 569.11 and AICcWt = 0.87, Tables 3 & 4, compared to next best model AICc = 574.91, and AICcWt = 0.05,). Hence, distance and the interaction term were found to be significant, while cat density and satiation index were not.

For analysis two, when variables were examined individually, cat density (marginal R^2^ = 0.33, conditional R^2^ = 0.83, p = 0.008, n = 96, Poisson GLMM, Fig 6) and cat satiation (marginal R^2^ = 0.39, conditional R^2^ = 0.82, p = 0.002, n = 96, Poisson GLMM, Fig 7) were significantly negatively associated with the number of reptiles observed in a survey. The full model that best fit the data included cat density (p = 0.047) cat satiation (p = 0.218), and the interaction between cat density and cat satiation (p = 0.001) (marginal R^2^ = 0.67, conditional R^2 =^ 0.83, AICc = 686.08 and AICcWt = 0.89, Tables 3 & 5, compared to the next best model AICc = 692.09 and AICWt = 0.04, Table 3). In this model, cat satiation was not significant, cat density was significant, and there was a significant negative interaction between cat density and cat satiation.

**Fig 6.**
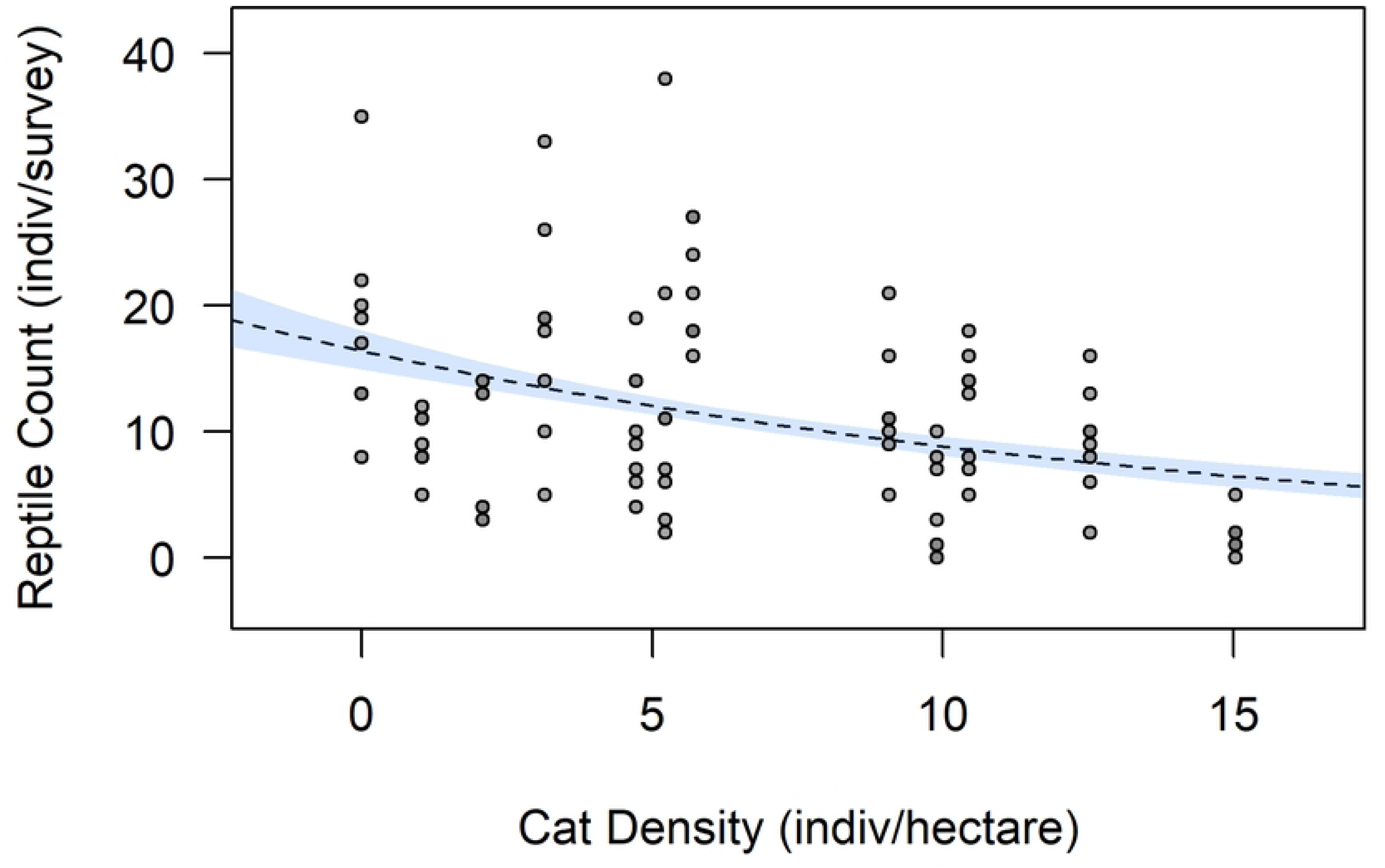
Poisson generalized linear model showing association between reptile counts per survey and cat density. Shaded region represents 95% confidence interval for fixed effects, p = 2e-16, null deviance = 544.02 on 95 degrees of freedom, residual deviance = 462.78 on 94 degrees of freedom.

**Fig 7.**
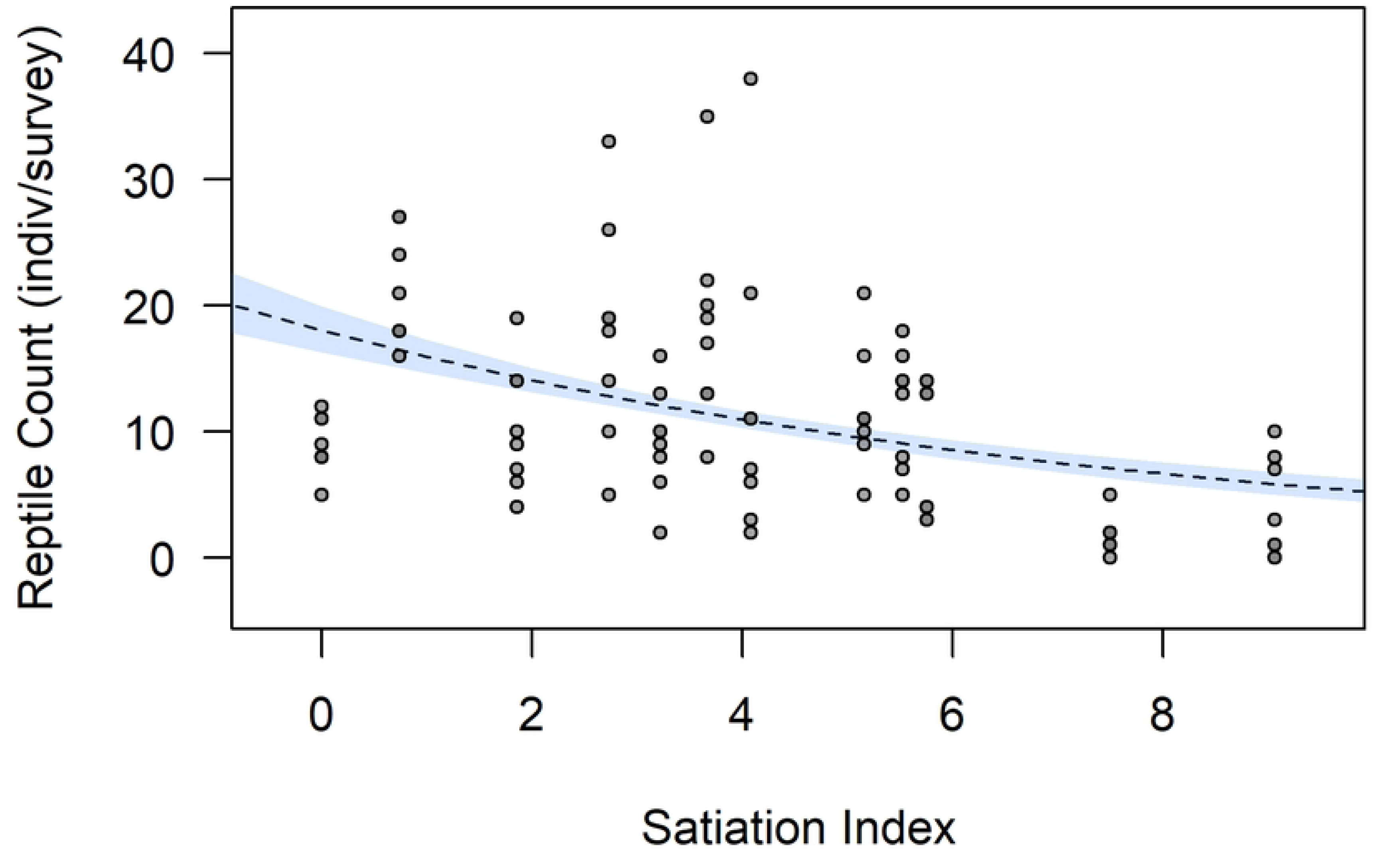
Poisson generalized linear model showing association between reptile counts per survey and satiation index. Shaded region represents 95% confidence interval for fixed effects, p = 2e-16, null deviance = 544.02 on 95 degrees of freedom, residual deviance = 440.90 on 94 degrees of freedom.

**Table 3:** Model selection for factors predicting number of reptile observations from two different analyses. Model A shows results from reptile analysis 1 and Model B shows results from reptile analysis 2. Models are ranked in ascending order by corrected Akaike corrected information criteria (AICc). ΔAICc, AICc weight, and Log Likelihood are provided for each model. Model variables include distance (model A only), cat density, cat satiation, and an interaction between cat distance and cat satiation.

|  | Reptile Count ~ | AICc | ΔAICc | AICcWt. | Log Likelihood |
| --- | --- | --- | --- | --- | --- |
| **A** | *Reptile Analysis 1* |  |  |  |  |
|  | ~ **distance + cat density + cat satiation + interaction** | **569.11** | **0.00** | **0.87** | **-278.09** |
|  | ~ distance + cat satiation | 574.91 | 5.79 | 0.05 | -283.23 |
|  | ~ distance + cat density + cat satiation | 575.04 | 5.93 | 0.04 | -282.19 |
|  | ~ cat density + distance | 576.16 | 7.05 | 0.03 | -283.86 |
|  | ~ cat density + cat satiation + interaction | 579.63 | 10.51 | 0.00 | -284.48 |
|  | ~ distance | 579.83 | 10.71 | 0.00 | -286.78 |
|  | ~ cat satiation | 585.52 | 16.4 | 0.00 | -289.63 |
|  | ~ cat density + cat satiation | 585.61 | 16.49 | 0.00 | -288.58 |
|  | ~ cat density | 586.77 | 17.66 | 0.00 | -290.26 |
|  | null model | 590.48 | 21.37 | 0.00 | -293.18 |
| **B** | *Reptile Analysis 2* |  |  |  |  |
|  | ~ **cat density + cat satiation + interaction** | **686.08** | **0.00** | **0.89** | **-337.71** |
|  | ~ cat satiation | 692.09 | 6.01 | 0.04 | -342.91 |
|  | ~ cat density + cat satiation | 692.24 | 6.16 | 0.04 | -341.90 |
|  | ~ cat density | 693.59 | 7.51 | 0.02 | -343.66 |
|  | null model | 697.27 | 11.2 | 0.00 | -346.57 |

### Heat maps

Heat maps showing the spatial distribution of cats, dumpsters, and food dishes were created for each village in ArcGIS Pro v3.4 to provide visual supplementation to these results. Heat maps include every cat observation across all three surveys in each village. These heat maps clearly show that cats form high density aggregations near human-derived food resources. Due to large cat population size, these results were best visually observed in the village of Vivlos (Fig 8), where the highest cat density regions (red regions) are all associated with a dumpster or a food dish, while none of the lowest cat density areas (blue regions) had a detectable source of human-derived food within them. Results from heat maps from other villages with smaller populations were consistent with these findings.

**Fig 8.**
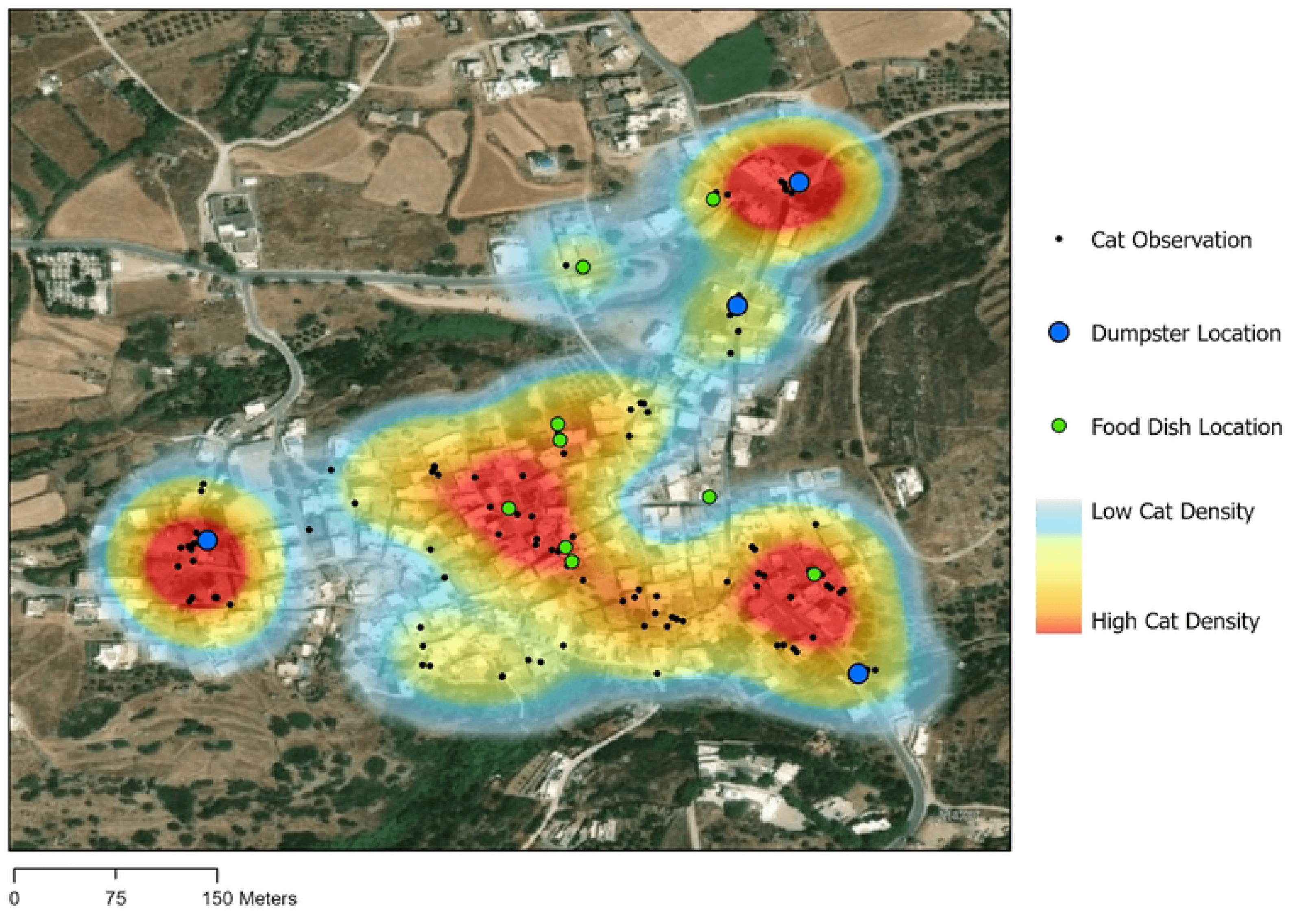
Heat map showing the spatial distribution of cats (n = 171 cat observations) in relationship to human subsidies in a typical Naxian village (Vivlos). Black dots show all cat observations made during three nights of surveys. Cats typically stayed in close proximity to sources of food, whether deeding dishes or dumpsters Map created in ArcGIS Pro v3.4. Produced with ESRIi World Image WGS1983 World Image Base Map (2025) [62].

## Discussion

Cats pose an increasingly pressing ecological problem across most island ecosystems globally [14–15], yet managers and local decisionmakers still have a poor understanding of the magnitude of the impacts, as well as the causative factors. By surveying outdoor cats along a gradient of increasing food supplementation at twelve discrete sites within a single representative Mediterranean ecosystem, this study demonstrated that cat population size is the direct product of the quantity of human-derived food resources available to them. It was found that food resources come primarily in the form of food scraps or dry cat food placed by local residents in their yards or village paths. Equally important are the communal dumpster bins that are usually open and sought out by groups of stray cats. These results demonstrate that cat densities in villages are correlated with the density of both these resources. Beyond density of food resources, a separate and distinct effect of human population on cat density was detected, potentially reflecting additional benefits in the form of shelter and medical care some cats derive from a subset of each community’s inhabitants. While previous studies have shown that cats will congregate and form high density colonies around easily accessible food resources [34,63,64], these results further demonstrate that cats are not just simply reorganizing themselves spatially around these resources (Figs 4 & 10), but rather that population level cat density readily rises to match bottom-up human supplementation effects.

These analyses assume that the cat population within each village constitute a discrete unit. The existing literature indicates that while the home range size of cats varies widely [64–65], home range decreases with increasing productivity of the landscape [66]. The high density of food dishes and garbage sites within each village in this study (Fig 8) indicates that sufficient resources are available to collapse home ranges and congregate cats within the individual village. The cat surveys, conducted over three consecutive nights demonstrate high level of philopatry of individual cats with most animals not moving more than a few dozen meters from day to day. The assumption of closed village populations was further supported by a conspicuous absence of cats away from human settlements (personal observation). These observations agree with past research [11] and indicate that cats are not roaming the landscape of Naxos or travelling between villages in significant numbers, but rather remain close to their within-village food sources. Consequently, increases in cat density within a village are the result of local legacies rather than the result of extensive inter-village movement.

Is it possible that the association between cats and food dishes is simply the result of increased provisioning by compassionate locals in response to rising cat populations, rather than the other way around? While some of this cannot be completely ruled out, field observations suggest instead that feeding intended for a local house pet ends up inadvertently supporting other stray cats and hence promotes population increases. Furthermore, cat populations closely reflect the density of garbage dumpsters, the number of which certainly is not determined by cat density. In summary, the existing evidence suggests that intentional or unintentional supplementation of cats drives population numbers rather than vice versa.

We found that cat population density is negatively correlated with resident reptile population abundance, indicating substantial levels of predation on lizards and snakes. This corroborates regular but *ad hoc* observations during the study of cats consuming wildlife (S1 Appendix). While data collection in this study focused on reptiles, parallel trends are known to occur within birds as well [40].

This study also reveals that while cat impacts on local wildlife are both pervasive and density-dependent, they are also modulated by the degree of cat satiation. Because cat populations do diverge somewhat from what would be predicted based on available resources, the residuals of this relationship were used as a metric of the relative resources available per cat capita, essentially obtaining an index of over/under provisioning (termed cat satiation). The results indicate that not only is the extent to which cat satiation is proportional to decreases in wildlife populations independent from cat density (Fig 7), but also that the interaction between cat density and satiation is particularly important. Hence, cat numbers are more negatively associated with wildlife populations when satiation indices are high, i.e. cats are hungry. This is consistent with a scenario where cats preferentially consume human-derived resources and switch to preying local wildlife when these are not sufficient. Despite the strong evidence of direct cat impact on reptiles through predation, a different explanatory process cannot be completely excluded. In this scenario, higher cat density and reduced provisioning results in reptiles that are more wary, more cryptic, and less active, therefore reducing detectability during surveys.

A clear linear rise in reptile abundance with increasing distance from a village edge was found (Fig 5). Reptile counts increased on average by ∼40% from the immediate edge of the village to 200 meters away from a village, suggesting that proximity to villages was detrimental to reptiles. While this pattern is consistent with increased cat-induced mortality close to villages, there was no relationship between cat density and how rapidly reptile numbers rose with increasing distance from a village. Hence, the steepness of this decline in reptile abundance when approaching a village was not significantly related to cat density or cat satiation, perhaps because of insufficient sample sizes or non-linear relationships. While the possibility that cats do not influence the effect of distance is not being ruled out, alternative causes may be that 1) as lizards are predated by cats, the high density of highly active lizards means that the new spaces created by cat predation are readily filled by a lizard from further away and 2) the increase in reptiles away from villages could be attributed to other factors such as habitat suitability increasing with greater distance from human settlements.

Because anthropogenic food subsidies are critical for cat population growth, and increasing cat population size has proportional negative impacts on resident wildlife population sizes, this study strongly indicates that human subsidies are facilitating hyperpredation of wildlife on Naxos. As a result, in many Mediterranean regions, human supplementation drives the impacts of an artificially elevated cat population on resident wildlife.

### Management recommendations

Public opinion regarding free-ranging cats on the Aegean islands varies greatly, with some residents considering them pests while others taking care of them through regular feeding (personal observation). On Naxos and other islands, cats are increasingly becoming a source of amusement for tourists, and cat-themed merchandise can be found on sale in many retail stores. Local businesses have also capitalized on the tourist’s affection for cats by selling bags of cat food to allow for a petting zoo-like experience with the local strays. Over the last several years, there has been a pronounced shift in public opinion regarding cat welfare away from the past *laissez-faire* attitude toward a more compassionate stance. As a matter of fact, it is the attitude of support and consequent feeding that has resulted in the present-day ballooning of cat populations on Aegean islands. Concurrently, there is a growing realization that cat overpopulation is a problem, primarily due to animal welfare rather than wildlife conservation reasons, and cat population control programs are increasingly discussed.

This study has shown that increasing cat densities translate directly into higher mortality for reptiles, and that the negative impacts of cats is strongest when human food becomes limiting, and cats are forced to rely more on wildlife for nutrition. These results may be interpreted as providing conflicting management suggestions. While in the short term, overprovisioning of cats reduces predation of wildlife, in the long term this will result in accelerated reproduction and expanding cat populations. This conflict between short-term benefits versus long-term detriments can best be navigated through a carefully balanced approach. If cat populations are to be managed as to reduce wildlife mortality, several steps need to be taken including expansion of existing trap-neuter-release programs, as well as reduction in supplemental feeding of unsterilized cats to reduce reproductive potential. More specifically, proposed recommendations include 1) improving waste management policies to decrease accessibility of this resource to cats, 2) increasing trap-neuter release efforts, and 3) providing local education about the deleterious impacts of cats on wildlife and encouraging both residents and tourists to only feed sterilized cats. This last point is significant, as keeping only sterilized cats fed will decrease their dependence on wildlife while also not contributing to cat population growth. The reduced access to human-derived food by unsterilized cats through recommendations 1 and 3 may cause them to consume wildlife more in the short term, but the loss of easily available human food resources should also reduce their reproductive output. While controversial, it is worth mentioning that humane cat culling is an option as well, as it has been shown to be highly effective, especially on islands [67]. Additionally, the implementation of cat management needs to be followed by close environmental monitoring to assess unintended consequences such as mesopredator release of rats following cat population declines [22].

### Conclusion and future research directions

As global biodiversity continues to be lost at an alarming rate [68], the importance of understanding the mechanisms driving species extinction is increasing. The conclusions of this study should be applicable to numerous regions in the Mediterranean Basin, at least those where cats impact small- and medium-bodied vertebrates. Further research efforts need to center on investigating how the extent of human subsidies influences that distance cats will roam from a village, and stable-isotope analysis of scat can help elucidate how the ratio of anthropogenic and natural food sources vary along this gradient. Lastly, this study focused on the high tourism summer period when large numbers of visitors come to the island. Extending the study to the winter months when less food supplementation is available can help to understand the true impact on resident wildlife.

## Acknowledgements

We would like to thank Father George Palamaris for providing us with housing in the Venetian castle of Naxos while conducting this research, and to Mayor Dimitris Lianos for being supportive of the research the Foufopoulos lab conducts in Greece. We would also like to thank Andressa Viol for her assistance in developing conceptual figure 1.

---

**S1 Table.** Summary table showing data used in this study. Area and density columns are in hectares, total food density column is equal to the summation of dish density and dumpster density, the reptile count column shows the average and standard deviation of total reptile counts observed at the corresponding village across all surveys.

**S1 Appendix.** A. Outdoor cats line a set of dumpsters in Chalkio. B. Outdoor cats feed out of a feeding dish in Vivlos. C. Cat predates a Eurasian Collared Dove in Chora. D. Cat predates a wall lizard in Moni.

**S2 Appendix.** Photo taken in Glynado demonstrating the exceptional densities cats can achieve in urban environments.

